# Multimodal microscale mechanical mapping of cancer cells in complex microenvironments

**DOI:** 10.1101/2022.03.28.486131

**Authors:** Miloš Nikolić, Giuliano Scarcelli, Kandice Tanner

**Affiliations:** Laboratory of Cell Biology, Center for Cancer Research, National Cancer Institute, National Institutes of Health, USA; Maryland Biophysics Program, IPST, University of Maryland, College Park, MD, USA; Fischell Department of Bioengineering, University of Maryland, College Park, MD, USA

**Keywords:** Microrheology, Physical properties, Optical methods, Cancer cells, Topographical cues, Engineered matrices

## Abstract

The mechanical phenotype of the cell is critical for survival following deformations due to confinement and fluid flow. One idea is that cancer cells are plastic and adopt different mechanical phenotypes under different geometries that aid in their survival. Thus, an attractive goal, is to disrupt the cancer cells’ ability to adopt multiple mechanical states. To begin to address this question, we aimed to quantify the diversity of these mechanical states using *in vitro* biomimetics to mimic *in vivo* 2D and 3D extracellular matrix environments. Here, we used two modalities Brillouin microscopy (∼GHz) and broadband frequency (3-15kHz) optical tweezer microrheology to measure microscale cell mechanics. We measured the response of intracellular mechanics of cancer cells cultured in 2D and 3D environments where we modified substrate stiffness, dimensionality (2D versus 3D), and presence of fibrillar topography. We determined that there was good agreement between two modalities despite the difference in timescale of the two measurements. These findings on cell mechanical phenotype in different environments confirm a correlation between modalities that employ different mechanisms at different temporal scales (Hz-kHz vs. GHz). We also determined that observed heterogeneity in cell shape that is more closely linked to the cells’ mechanical state. We also determined that individual cells in multicellular spheroids exhibit a lower degree of mechanical heterogeneity when compared to single cells cultured in monodisperse 3D cultures. Moreover, the observed decreased heterogeneity among cells in spheroids suggested that there is mechanical cooperativity between cells that make up a single spheroid.

## Introduction

Cancer cells encounter many different biochemical and physical cues within the organ microenvironment. These cues are anisotropic and undergo temporal evolution within the microenvironment milieu [1]. As part of an intricate feedback mechanism, cancer cells sense environmental cues, which evokes a modulation in cellular behavior, which in turn regulates secretion of extracellular cytokines, alterations in tissue architecture and remodeling of cell mechanics [1, 2]. Thus, there is an emergence of distinct microenvironments during cancer progression. Simply, the cues at a stage of transition from normal to malignant may be distinct from those present at stages of invasive growth into surrounding tissues due to the continuous tissue remodeling. [2, 3]. Understanding the interplay between dynamic tissue reciprocity and the emergence of heterogeneous cell phenotypes is critical for our understanding of why some cancer cells remain indolent and why some are aggressively invasive.

This dynamic reciprocal cross talk may also drive phenotypic changes that might select for clones with enhanced survival, motility, and drug resistance [4-8]. These phenotypes can be classified by genetic, metabolic, and physical traits [9, 10]. The latter is often referred to as the mechanical phenotype. The cellular constituents spanning a multiplicity of length scales, enzymatic activity, cell cycle stage collectively regulate the physical phenotype of the cell [11]. In the last decade, mechanical phenotypes have been posited as critical determinants of cancer progression [12-18]. Specifically, the mechanical phenotype can regulate a cell’s response to external forces such as those encountered during invasion and transit within conduits during circulation [19, 20]. Mechanical phenotype also is an important determinant of motility strategies such as amoeboid, mesenchymal and “piston” driven modes of migration that are needed to navigate complex 3D structures [21, 22]. Finally, mechanical phenotype also influences the homotypic and heterotypic cell-cell interactions that occur within the tumor and interactions with stromal and immune cells [23, 24]. These cellular couplings such as cancer cell-cancer associated fibroblasts, cancer cell-cancer associated macrophages and clusters of cancer cells have all been shown to facilitate tumor outgrowth, escape, invasion, extravasation, and colonization of distant organs [25]. Our understanding of the environmental regulation of these states necessitates technical expertise to resolve mechanical phenotypes in complex environments.

The mechanical phenotype can be defined by several metrics such as a viscoelasticity, cell shape, cell deformability and adhesion properties. Of these metrics, single cell viscoelasticity provides a metric to assess the material properties of individual cells. Microrheology is a common metric that is used to quantify material properties [26-28]. These material properties are defined by combinatorial contributions due to cellular components that show energy dissipation and elastic properties. Novel tools such as atomic force microscopy, optical tweezers, magnetic twisting cytometry and micropipette aspiration have been utilized for characterization of cellular material properties [10, 29, 30]. However, it remains technically difficult to assess cells in complex environments with microscale resolution. To address this technical need, we methodically probed the single cell mechanics for a range of *in vitro* assays as a function of dimension, anisotropy and multicellular organization using two optical based techniques that can access the mechanical properties of cells in complex 3D environments. We employed Brillouin microscopy [31, 32] which is sensitive to material properties at the GHz timescale and broadband frequency multiplexed optical tweezer microrheology (3-15kHz) [33, 34]. These techniques allowed us to probe microscale mechanical properties and serve as a platform for comparative studies at the same length scale, in different frequency regimes.

## Materials and Methods

### Cell culture

Human glioblastoma cells - U87(ATCC® HTB14-™) were obtained from ATCC, MCF10CA1h cells [35] were received from the Barbara Ann Karmanos Cancer Institute (Detroit, MI, USA), and they were cultured as previously described. Briefly, the U87 cells were cultured in DMEM – Dulbecco’s Modified Eagle Medium (ThermoFisher Scientific, 11995065) that was supplemented with 10% fetal bovine serum (FBS) (ThermoFisher Scientific, 11995065) and 50 U/mL penicillin and 50 µg/ml streptomycin (Thermo Fisher, 15070-063). MCF10CA1h cells were cultured in the complete medium: DMEM/F12 (11330-032, Thermo Fisher Scientific), 5% horse serum (16050-122, Thermo Fisher Scientific), 5 ng/ml EGF (AF-100-15-1MG, Peprotech), 0.5 mg/ml Hydrocortisone (H0888-1G, Sigma-Aldrich), 100 ng/ml Cholera toxin (C8052-2mg, Sigma-Aldrich), 10 µg/ml insulin (I1882-100MG, Sigma-Aldrich) and 1x penicillin/streptomycin solution (15070-063, Thermo Fisher Scientific). Cells were cultured at 37°C, in 5% CO_2_. Cell propagation was performed by detaching adherent cells using Trypsin (0.25% for U87 cells (ThermoFisher Scientific, 80-2101) and 0.05% for MCF10CA1h cells (25-052-Cl, Corning)) as previously described. All experiments were performed using cells with passage number less than 19. Cell medium was changed every 2-3 days.

### Preparation of polyacrylamide substrates

Polyacrylamide gels of varying stiffness were fabricated using previously reported protocols [36]. Briefly, 35mm glass bottom dishes (Ibidi, 81158) were incubated with 0.1 M sodium hydroxide (Sigma-Aldrich, 72068). Then, 200 µl APTMS was added to the dishes for 3 min (Sigma-Aldrich, 281778), followed by 400 µl of 0.5% glutaraldehyde for 30 min (Sigma-Aldrich, G6257) to ensure polyacrylamide attachment to the treated glass. The stiffness of polyacrylamide gels can be altered by tuning the relative ratios of acrylamide to bis-acrylamide. First, a mixture of 40% acrylamide (Sigma-Aldrich, A4058-100ml), 2% bis-acrylamide (Sigma-Aldrich M1533-25ml), phosphate buffered saline (PBS) (ThermoFisher Scientific, 14190-144), TEMED (Sigma-Aldrich, T7024-25ml), and 10% w/v ammonium persulfate (Sigma-Aldrich, 215589) were thoroughly mixed. In these experiments, we fabricated gels that corresponded to a shear modulus (G’) of 0.1 kPa (soft gels) and 32 kPa (stiff gels). For soft gels, the final concentrations were 5% acrylamide, and 0.04% bis-acrylamide. For stiff gels, the final solution contained 18% acrylamide, and 0.4% bis-acrylamide. In each case, a total volume of 500 µl was prepared. 30 µl of the gel solution was added to the pre-treated 35mm glass bottom dish and covered with a round coverslip 18 mm round coverslips (#1, thickness) to create a circular gel. This coverslip was pre-treated with RainX (Illinois Tool Works) for 5 minutes to make them less adhesive to the polyacrylamide gels. After 15 min, 2 ml of PBS was added to the dish. After an additional 15 min, the glass coverslip was gently dislodged using tweezers, leaving a polyacrylamide gel ∼100 µm in thickness.

Cells do not attach to biologically inert polyacrylamide and require coating the gel surface with an extracellular matrix protein to promote attachment. First, the polyacrylamide gels were treated with 0.25 mg/ml Sulfo-SANPAH (ThermoFisher) in 50 mM HEPES (ThermoFisher). They were coated with 200 µl of the sulfo-SANPAH solution and placed under the UV lamp (UVP, Blak-Ray B100AP, 100W, 365nm) for 6 min. This step was repeated twice. The gels were then gently washed with 50 mM HEPES buffer twice and covered with a solution of fibronectin (Milipore Sigma FN010) - 10 µg/ml in HEPES for 12 hours at 4°C. Afterwards the gels were washed twice with PBS. Next, 2 ml solution of 50,000 cells in the serum free medium was added to the gels. Gels were then placed in the incubator overnight to facilitate cell attachment.

### Preparation of cells on 2D and in 3D culture

Cells cultured on top (2D) and embedded in 3D laminin rich extracellular matrix were prepared as previously described [37]. Briefly, we coated the bottom of a chilled 4 well µ-Slide (Ibidi, 80427) with 100 µl of ice-cold Matrigel (Corning, 356231, lot 8232015). The slide was then incubated at 37°C for 5 minutes for the Matrigel layer to solidify. For seeding cells in 2D, 50,000 cells were added directly to the well in 500 µl of serum free medium. For seeding cells in 3D, 430 µl Matrigel was mixed on ice with 50 µl of 2 × 10^6^ cells/ml solution of cells (10^5^ cells total) in serum free medium, and 12 µl of serum free cell medium. The mixture was added on top of the previously gelled layer of Matrigel and incubated at 37°C for 30 minutes for complete gelation. After 30 minutes, 350 µl of serum free medium was added each well. Cells were used imaged the following day.

### Preparation of fibrillar topography (FT) in 3D culture

Aligned magnetic self-assembled fibrillar matrices were fabricated using a previously reported protocol [37-39]. First, human fibronectin (Milipore Sigma, FN010) was fluorescently labelled using the DyLight™ 488 Microscale Labeling Kit (Thermo Scientific, 53024) according to the supplier protocol. Concentration of the fluorescently labeled protein was measured using a spectrophotometer (Nanodrop 2000c) and kept for up to 4 weeks at 4°C. Fluorescently labelled human fibronectin was conjugated to the carboxylated magnetic polystyrene beads (Ademtech Carboxy-Adembeads Coupling Kit, 02820) according to the manufacturer protocol. Magnetic beads, 300 nm diameter were washed three times in the activation buffer. The beads were resuspended at the concentration of 0.5mg/100µl. Activation of the beads was achieved by incubation in a 2 mg/ml solution of EDC at room temperature for one hour. Then, 20 µg of fluorescently labelled protein was added to 100 µl of the activated beads and incubated at room temperature overnight under gentle shaking. The next day, the conjugated beads were washed three times with storage buffer and kept at 4°C for up to 1 week at concentration of 10 mg/ml.

The fibrillar topography in 3D (FT) sample was prepared similarly to the 3D Matrigel encapsulation of cells as described above. 430 µl of ice-cold Matrigel was mixed with 12 µl of FN-conjugated beads (10 mg/ml), and 50 µl of the cell solution (total 10^5^ cells in serum free medium) and added to the well of the 4-well Ibidi slide that had a layer of 100 µl of Matrigel previously polymerized on the bottom. To induce alignment, the slide was then placed on ice-cold magnet (KJ Magnetics, B×8×8×8-N52) for 15 minutes. After 15 minutes, the beads assemble in a fibrillar topography. Sample was then immediately placed in the incubator at 37°C for 30 minutes for Matrigel to solidify. Afterwards 350 µl of serum free medium was added to the well. A control sample (FT control) was prepared in the same way, but incubated on ice for 15 minutes, in a separate ice bucket. The FN-conjugated magnetic beads were uniformly dispersed through the 3D gel for the FT control sample.

### Sample preparation for optical tweezer experiments

Cells were detached from the culture flask using 10mM EDTA solution. Cells were resuspended in fresh medium at concentration of 1.5 ×10^6^ cells/ml. Cells were then mixed with a solution of polystyrene beads (1 µm diameter, 2% solids, Invitrogen, F8816) in proportion of 100 µl beads/500µl cell solution, and incubated at 37°C, 5% CO_2_ for 30 minutes with gentle mixing for 30-45 min at 37ºC. This step resulted in internalization of beads by the cell through the process of phagocytosis. The cells were then centrifuged at 150g for 5 min (same as during cell passaging) and resuspended in PBS to remove excess beads. They were centrifuged and resuspended once more to obtain the needed concentration (2 × 10^6^ cells/ml) in the serum free medium. This suspension of cells with internalized beads was then used for preparing the 2D and 3D culture samples as described above.

### Spheroid morphogenesis assay in 3D on-top Matrigel culture

We followed established protocols to grow tumor-like spheroids of MCF10CA1h cells [40, 41]. Briefly, to create spheroids from cells seeded in the on-top configuration of, first 200 µl of ice-cold Matrigel (Corning, 356231) was added to a chilled 2-well imaging slide (Ibidi, 80287). Matrigel was spread evenly on the bottom of the glass using the pipette tip. The imaging slide was placed in the incubator for 30 min for the Matrigel layer to solidify. Cells were detached from the T25 flask as described above and suspended in fresh assay medium. Assay medium is the low-serum version of the complete medium: containing only 2% horse serum, while all the other ingredients are the same. Cells in suspension were mixed thoroughly with a pipette to ensure single cell suspension. Cell concentration was estimated using a hemocytometer, and 20,000 cells were added to the 500 µl of fresh assay medium. This cell solution was slowly and evenly pipetted into the 2-well on top of the solid Matrigel bed. The imaging slide was placed in the incubator for 30 min during which the cells settled on top of the Matrigel layer. Next, another 500 µl of assay medium containing 10% of Matrigel was added on top to create final concentration of 5% Matrigel in the well. Cells form in colonies over several days. Every 2 days the medium in the well was carefully aspirated and replaced with fresh 5% Matrigel containing assay medium.

### Brillouin Microscopy

Brillouin microscopy is a spectroscopic technique that measures the inelastic scattering of light from thermal density fluctuations inside the material (spontaneous Brillouin scattering) [31, 42, 43]. The scattered light undergoes a characteristic frequency shift when it scatters from these thermal phonons, and this frequency shift depends on the index of refraction n, mass density ρ and the longitudinal elastic modulus *M*^*′*^ of the probed material:

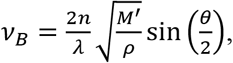

where *ν*_*B*_ is the measured Brillouin shift, and θ is the scattering angle an experimental constant (180° in our experimental setup). Longitudinal modulus M’ is the elastic constant defined as the constant that relates the uniaxial stress and uniaxial strain during compression. Therefore, Brillouin shift is an all-optical and high resolution measurement of local mechanical properties of a given material [31]. Following the current practice, in this study we report the value of Brillouin shift in GHz as a proxy measurement of the mechanical properties of live cells, with the assumption that the 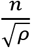 factor does not vary significantly in biological materials [44-46].

#### Brillouin imaging

Brillouin microscopy of cells was performed as described before [32, 44]. We used 60x, 0.7 NA objective (Olympus LUCPLFLN60X) to illuminate the sample with a light from a 660 nm laser (20-30 mW). The backscattered light was coupled into a single mode fiber and analyzed using the two VIPA cross-axis spectrometer with additional spectral purification elements (apodization and coronography [32]). Each pixel in Brillouin images comes from one acquired Brillouin spectrum (Figure 1 A). In the VIPA spectrometer the frequencies of the light are separated in space and imaged on a high sensitivity EMCCD camera (Andor iXon 897). Stray light from elastic scattering was blocked using adjustable slits in the spectrometer. Region that contains anti-Stokes Brillouin scattering peak and Stokes peak of the next diffraction order was collected. 5 pixels were averaged in the direction perpendicular to the spectral dispersion axis to obtain the intensity vs. frequency (in pixels) graph that contains the two Brillouin peaks (Anti-Stokes and Stokes of next diffraction order). Each of the two peaks was fitted with a Lorentzian function in a custom MATLAB program using nonlinear least squares fitting to localize peak centers. The distance between two peaks was calculated and used in further analysis to remove effects of laser frequency drift throughout the experiment. To calibrate the spectrometer, we measured the average of 500 Brillouin spectra of water and methanol collected with exposure time of 10 ms. By using the known literature values of Brillouin shift of water and methanol the spectral dispersion parameter (GHz per pixel) and the effective free spectral range (FSR) of the spectrometer were calculated. Calibration was performed at least once and hour and after each experimental condition. Exposure time (pixel dwell time) used in cell imaging was in the range 20 and 50 ms, depending on the imaging depth. Typically, each measurement has shift precision of approximately 8 MHz (0.13%). Samples were placed on a 3D motorized stage and scanned across the stationary laser focus. For each imaged cell, we mapped Brillouin shift in a single horizontal or vertical plane passing through the middle of the cell.

**Figure 1.**
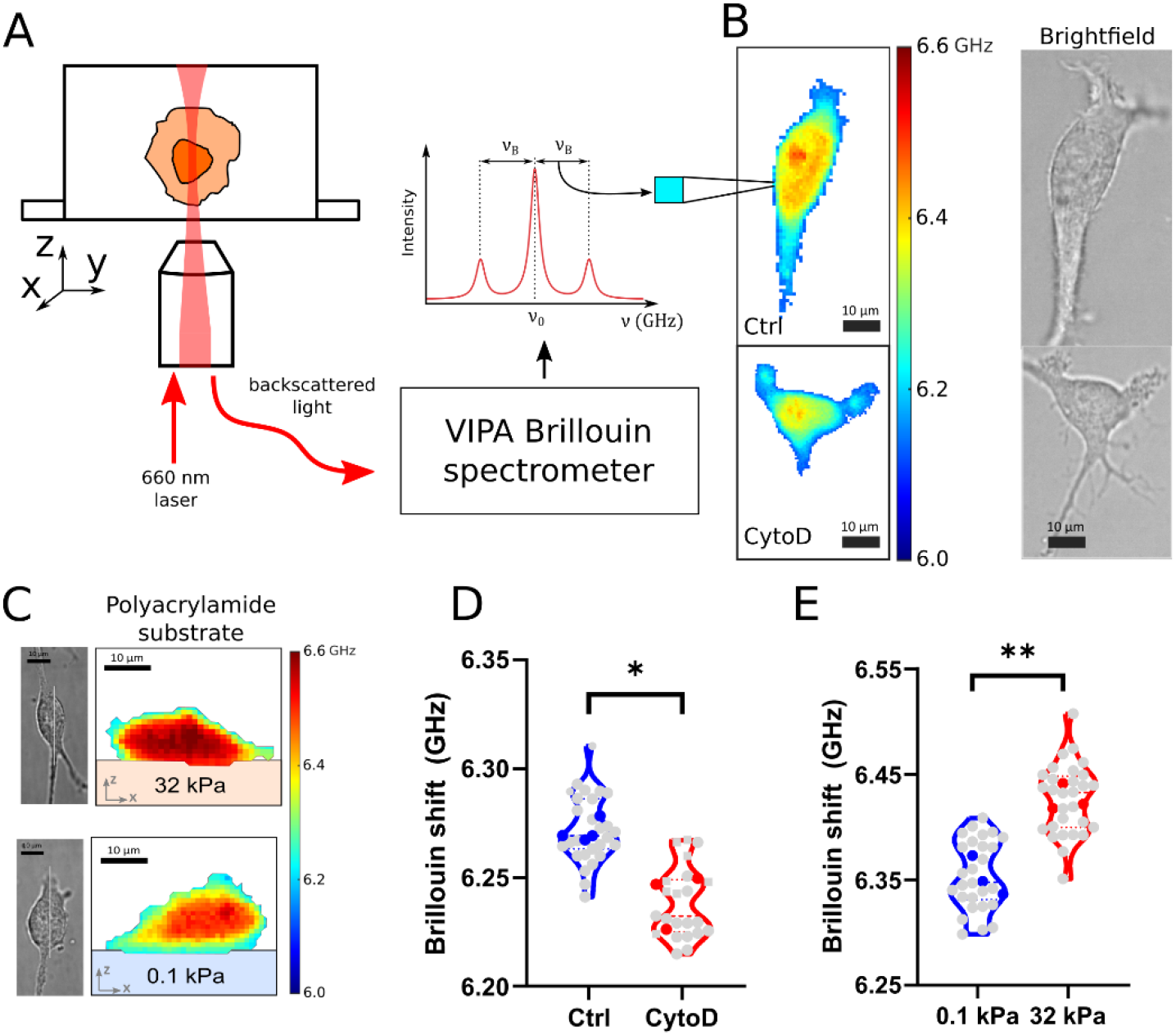
(A) Schematic of the experimental setup. Cells are prepared in an *in vitro* microenvironment (2D or 3D). Cells are illuminated with a 660 nm laser, and backscattered light from the confocal volume is analyzed using the VIPA Brillouin spectrometer. For each pixel of the image, the spectrum is analyzed, and Brillouin frequency shift is recorded. (B) Brillouin shift maps and brightfield images of live U87 cells: control and treated with cytochalasin D. Images represent a horizontal x-y confocal slice. Right panels are brightfield images of the same cells. (C). Brillouin shift maps and brightfield images of live U87 cells cultured on polyacrylamide gels of 0.1 kPa and 32 kPa stiffness. Images represent the vertical x-z confocal slice of the cell whose location is denoted by the white line in the brightfield images. (C) Average Brillouin shift of cells treated with cytochalasin D (red) with respect to the control (blue) in three independent experiments. Each gray point represents average shift of all pixels in one cell, each color point is the average all cells in one experiment. (D) Average Brillouin shift of cells cultured on top of polyacrylamide gels of different stiffness in the same plot as in (C). Unpaired t-test was performed on the average values of independent experiments.

#### Brillouin image analysis protocol

The culture media and Matrigel show a lower shift than cells. We identified the cell in Brillouin maps by separating it from the shift of the culture media. We did this by manually selecting a set of pixels containing the surrounding medium and keeping only pixels that at least 3 standard deviations larger than the average value of Brillouin shift of the medium. However, for conditions where cells are cultured on polyacrylamide gels and in gels with fibronectin coated nanoparticle fibrils, the boundary between the cell and the polyacrylamide gels or fibrils was manually selected in the Brillouin shift maps using the *imfreehand* function in MATLAB. In these cases, a vertical slice is always mapped through the cell, to differentiate the gel and the fibrils in the Brillouin images. Cells cultured on top of polyacrylamide gels were imaged inside a microscope stage incubation chamber at 37°C, 5% CO_2_. The rest of the cells were imaged at room temperature conditions. Measurement of each sample was done within 1 hour of taking the cells out of the incubator.

In the case of measuring suspended cells, we added suspended cells to the glass bottom imaging dish immediately after harvesting from a T25 flask. We allowed 2-3 minutes for the cells to settle down on the glass bottom of the imaging dish. We then imaged large horizontal regions (∼ 150 µm by 150 µm) that include many cells. All measurements were performed within 20 min of adding cells to the dish, before cell attachment occurs [47]. We confirmed that within this time cell height does not change significantly by imaging vertical xz Brillouin shift slices through cells. To speed up the imaging process we also sampled the images with a larger 2 µm step size. Cells were identified in these Brillouin shift images by thresholding at a fixed threshold of 6.15 GHz (approximately >3σ from the value of Brillouin shift of the medium). Images were further transformed using watershed transform to separate cells from the background and to split isolated objects into individual cells (Figure 4. A). This automated image processing allowed a higher throughput measurement with around 60 suspended cells per dish.

In the case of Brillouin imaging of spheroids, one horizontal slice was mapped approximately through the middle of the spheroid. Individual cell masks were manually selected in each Brillouin image (Figure 4 D) in MATLAB. Centroid of each selected cell region was recorded, as well as the average Brillouin shift in that region. We quantified the mechanical variability of cells within a spheroid as a standard deviation of the Brillouin shifts of all selected cell within a spheroid. For each spheroid measured, we also calculated the Brillouin shift difference between each pair of cells. To create the plot in Figure 4 F, we plotted those differences as a function of the distance between cell centroid locations in the Brillouin images.

### Optical Tweezer based active microrheology

Optical tweezer experiments were performed on the experimental setup as described previously [33, 34]. Briefly, the experimental setup consists of a detection part (detection laser 975 nm and quadrant photodetector QPD), and trapping part (trapping laser 1064 nm, 100 mW power; 2D acousto-optic deflector (AOD); piezoelectric translation stage).

#### Sample Measurement

The condenser of the microscope was placed in the Kohler illumination and the sample was put in focus. The bead of interest is positioned precisely in the center of the optical trap. This is achieved by optically scanning across the bead in 3 directions (x, y, and z) using a piezo XYZ nano-positioning stage (Prior, 77011201) and recording the voltages on the detection path QPD. Specific relation, between voltage and nm-position β was measured *in situ* by fitting the central linear region of the detector response to bead position. This allows for precise localization of the bead during trap oscillation, as previously described [49]. To perform active microrheology measurements on the correctly positioned bead, the trap beam is oscillated while recording both the optical trap QPD signal (force) and the detection QPD signal (position). The oscillation of the trap beam is multiplexed as a superposition of sine waves of differing phase and frequency with the same amplitude (25.4 nm). Twenty frequencies are deliberately chosen as co-prime numbers in the range of 3Hz-15kHz, to avoid interference and to allow for multiplexed measurement at each frequency. The waveform is pulsed for 2 s, followed by 2s with the trap stationary. and the sequence is repeated 7 times. Before collecting the active microrheology measurement a 10 second passive bead spectrum is recorded. Instrument control and data acquisition are performed using custom programs (National Instruments, LabVIEW).

#### Estimation of the local complex shear elastic modulus

Before measuring the viscoelastic constant at the location of each bead, the optical trap stiffness is calibrated in situ based on the combination of the passive and active power spectra of the bead position [50, 51]. Viscoelastic response of the material in the linear regime at the bead location is modeled by a generalized Langevin equation with additional terms: a harmonic term for the applied optical trap force and memory terms for viscoelastic friction *γ*_1,*U*_ and hydrodynamic memory effects *γ*_2,*U*_ [51-54]. The equation of motion of an undriven bead is given by

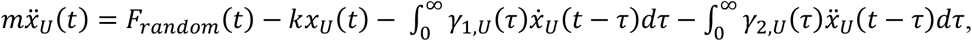

where, 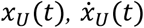, and 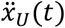 are the undriven bead position, velocity, and acceleration respectively; t is time, τ is correlation time, *F*_*random*_*(t)* is the Langevin noise term, *k* is the stiffness of the optical trap, *m* is the mass of the trapped bead, and *γ*_*1,U*_*(τ)* and *γ*_*2,U*_*(τ)* are the time-dependent memory functions representing friction and hydrodynamic memory effects respectively. The equation of motion for a bead driven by moving the position *x*_*L*_(*t*) of the laser is given by:

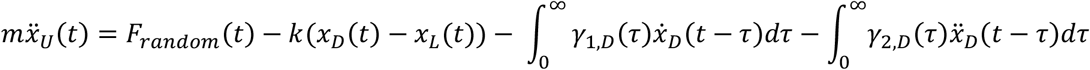

where 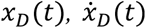, and 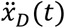 the driven bead position, velocity, and acceleration respectively; and *γ*_*1,D*_*(τ)* and *γ*_*2,D*_*(τ)* are the time-dependent memory functions representing friction and hydrodynamic memory effects respectively in the driven case. Onsager’s regression hypothesis which is a consequence of the fluctuation-dissipation theorem allows us to assume that the friction relaxation spectrum is equal in the driven and undriven system *γ*_*1,U*_*(τ)* = *γ*_*1,D*_*(τ)* and *γ*_*2,U*_*(τ)* = *γ*_*2,D*_*(τ)* [50-54]. This is valid in the case when the driven motion of the bead is on the same scale as the passive motion of the bead due to the Brownian motion. The spectrum of the actively driven bead is defined as 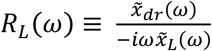,where 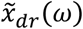 and 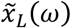 are the Fourier transforms of the positions as a function of time of the trapped bead and the trapping laser, respectively, that are recorded while the trap is oscillating. The active driven spectrum *R*_*L*_(*ω*) can be used to estimate the friction relaxation function 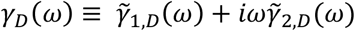 according to the following equation

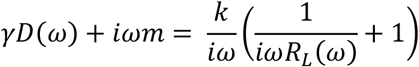

where ω is the trap oscillation frequency in rad/s, and ∼ indicates Fourier transform. The optical trap stiffness can be determined from the real part of the active power spectrum and the passive power spectrum and is given by 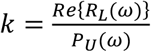, where *P*_*U*_(*ω*) = ⟨|*x*_*U*_ (*ω*)|^2^⟩, and 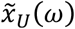 is the Fourier transform of the position of the of the undriven bead’s as a function of time while the trapping laser is stationary. The friction relaxation spectrum *γ*_*D*_(*ω*) is related to the complex viscoelastic modulus of the surrounding microenvironemnt by the generalized Stokes-Einstein relation 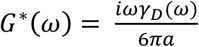, where *m* is bead mass, and *a* is the hydrodynamic radius of the bead. The complex viscoelastic modulus can be written as

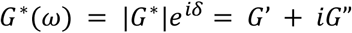

where |*G\**| *= (G’*^*2*^*+G”*^*2*^)^*1/2*^ is the magnitude and *δ* is the loss tangent *tan(δ) =* ^*G”*^*/*_*G’*_ and they encode rigidity and hysteresivity, respectively. The real part *G’* represents the storage (elastic) modulus and the imaginary part *G”* represents the loss (viscous) modulus.

#### Data analysis and statistics

For each cell analyzed, 3-5 beads were measured at different locations within the cell. Only cells exceeding ∼30 µm from the coverslip were analyzed in accordance with Faxen’s law. Between 7 and 10 cells were analyzed per sample in each experiment, and three independent experiments were performed (for 3 independent samples fabricated across different days and across different cell passage numbers). Data were analyzed using custom MATLAB and GraphPad Prism programs.

The measurements of the modulus magnitudes |*G*(ω)*|, *G’*, and *G”* for each single bead are distributed according to the log-normal distribution. The mean value of these magnitudes from the repeated measurements on the same bead are calculated using the maximum-likelihood estimate for the mean of the log-normal distribution [33].

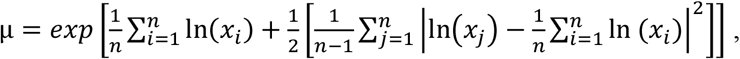

The maximum-likelihood estimate of the log-transformed variance is given by

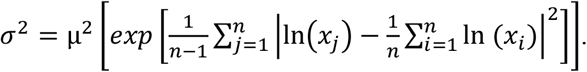

In case of the loss tangent which is normally distributed we use arithmetic mean and standard deviation as estimates of central tendency and dispersion among measurements.

We assume beads measurements within each cell and cells within the cell population are normally distributed. Active microrheology data are presented as mean complex, storage, or loss moduli vs frequency from 7 Hz to 15 kHz of all beads in a single cell. Average values of the G’ and G’’ for all cells in one dish (one experiment) were collected from three independent experiments. These (n=3) values were compared across all frequencies and presence of statistically detectable differences was analyzed with a two-way ANOVA test with Tukey’s post hoc correction.

### Cell shape analysis

Brightfield images of each cell sampled by Brillouin microscope was used to identify cell shape. Cell shape was detected using a custom-built MATLAB semi-automatic cell detection GUI program. Each image intensity was normalized and images were sent through an edge detection protocol [48] that created a binary gradient mask based on the Sobel algorithm. This binary image was dilated to connect edges, holes were filled, and the binary image was eroded (to undo the dilation while preserving the large objects – cells). Each detected preliminary cell shape was inspected and manually edited to select the correct cell region in the field of view using MATLAB’s imfreehand function. The edge detection protocol depends on the contrast of the brightfield images, and it is not always perfectly accurate, especially when there are multiple objects in the image in addition to the cell – e.g., aligned fibrils. For this reason, we manually inspected and corrected each image. Binary images of cells were analyzed in MATLAB to extract the relevant shape parameters: area, major and minor axis of fitted ellipse, and perimeter. We quantified the **aspect ratio** as the ratio of minor to major axis of the fitted ellipse, and **circularity** as perimeter^2^/(4π × area).

## Results

### Brillouin shift correlates with modulation of cell mechanics in cells cultured on 2D substrates

The actin cytoskeleton has been shown to be a key regulator of the mechanical properties of cells. We first assessed the Brillouin shift of cancer cells cultured in 2D in the presence and absence of a cytochalasin D, pharmacological inhibitor of actin [44, 55-58]. We determined that the average Brillouin shift of cancer cells decreases by 30 MHz (Fig. 1 A, C) when treated with 1 µM cytochalasin D for 30 min. Cells can modulate their internal mechanical properties, morphology, and organization of the cytoskeleton in response to substrate stiffness [59-61]. We next asked if there are differences in Brillouin shifts for cells cultured on different substrate stiffnesses. We determined that there is a distinct difference in Brillouin shift for cells grown on soft versus stiff substrates. Cells cultured on hard (32 kPa) polyacrylamide gels adopted Brillouin shift 75 MHz higher than cells cultured on soft polyacrylamide gels (0.1 kPa). We calculated the average Brillouin shift of a whole cell (gray points in Figure 1 D and E) and used those to estimate the mean Brillouin shift of cells in each condition (color points in Figure 1 D and E, one for each independent experiment).

### Cells cultured on top of hydrogels and embedded in hydrogel share similar mechanical phenotype

The effects of substrates with different properties (e.g. stiffness or ligand density [62]) on cell have been extensively studied. However, the effects of distance and boundary conditions on cells’ ability to sense a physical cue such as substrate stiffness is less understood. Specifically, if a cell is plated on top of a soft flat hydrogel (∼100 Pa) of a given thickness on a glass dish surface (>GPa), do cells behave comparably to a boundary condition where the soft substrate is infinite and isotropic in every direction? In other words, what is the boundary condition that governs mechanical phenotype of cells. We take advantage of the ability of Brillouin microscopy to non-invasively map mechanical properties of cells in 3D to address this question. We measured live cells embedded in 3D laminin rich hydrogel (Matrigel) and compared them to the cells grown on top of a flat hydrogel of the same composition. Cell morphological analysis using brightfield images revealed a diversity of shapes where some cancer cells in 3D were spherical and some adopted elongated shapes. Comparative analysis for cells grown on a 2D laminin rich ECM hydrogel show the characteristic elongated and flat morphology. Even though cells respond differently to 3D and 2D configurations, we consistently find no difference in Brillouin shift of cells in these two conditions.

To independently investigate the mechanics of cells in these two configurations, we performed the same experiments using high-frequency optical tweezer microrheology. This technique can probe complex elastic modulus of live cells directly in three dimensions by actively driving 1 µm beads embedded in the cell cytoplasm across frequencies ranging from 7 Hz to 15 kHz [33, 34]. (See Materials and Methods.) We again determined that the complex elastic modulus of cells cultured in in 3D and 2D configurations were comparable across all measured frequencies. The measured shear modulus at 19 Hz was G’_(3D)_= 39±11 Pa, and loss modulus G’’_(3D)_ = 25±7 Pa.

Fibrillar structures are present *in vivo* and alter cancer cell migration [63]. Cells can sense the fibrillar topography in their environment independently of the adhesion ligands that the fibrils present to the cell [37]. Using Brillouin microscopy, we set out to investigate whether cells also alter their mechanics in the presence of the fibrillar topography in 3D. Here, we used a model previously developed in our lab to fabricate fibrillar architecture in 3D [37-39]. We used magnetic nanoparticles coated with human fibronectin that in presence of magnetic field align into 1-2 µm thick fibrils that are embedded in the 3D hydrogel (Matrigel) together with cells. We previously determined that the overall mechanical and diffusive properties of the hydrogel were not altered due to the presence of the aligned particles. Instead, the presence or absence of cell protrusions were dictated by the aligned fibrils [37]. As mechanical phenotypes of cells are responsive to tissue anisotropy, we next asked if the mechanical phenotype is also altered. We determined that the mechanical phenotype of cells cultured in presence of fibrillar architecture and control conditions was similar as measured by quantifying the Brillouin shift.

### Cell mechanical state is a variable parameter that correlates to cell morphology

Within a given sample, cells adopt morphogenetic heterogeneities [64, 65]. We next asked if we can further identify sub types of mechanical phenotypes based on classification of different morphologies. First, we used the brightfield images of all cells which we imaged with Brillouin microscopy to determine cell morphology. To remove the underlying variability in mechanical properties of cell subpopulations in the data, we devised a classification scheme based on the circularity and aspect ratio of cell shapes. Circularity is defined as the ratio of the shape area and the square of its perimeter. It is normalized in such a way to be in the range between 0 and 1, where perfect circle has circularity of 1. Similarly, the aspect ratio is a measure of deviation from the perfectly round morphology. Here we define as the aspect ratio as the ratio of minor to major axis of the fitted ellipse to the shape boundary. This ensures that the aspect ratio also falls in the range from 0 to 1. By combining these two parameters, we can identify cells that either have comparable values of aspect ratio and circularity, or cells that have low circularity but intermediate or high value of aspect ratio. Within the first subset of cells that have comparable circularity and aspect ratio, we further divide those into two groups: round cells and elongated but not protrusive (type A: round cells AR>0.75, and type B: elongated cells AR<0.75). The remaining subset of shapes with very low circularity for the given aspect ratio (circularity – AR > 0.2) revealed cells of very long perimeter which indicated morphologies with an abundance of protrusions (Type C).

Using this classification, we identified detectable differences in Brillouin shift among these three shape types (Fig 3. C). We then focused on one morphological classification and asked if culture conditions then determined the Brillouin shift. We determined for the type B cells – elongated but not protrusive – adopted a higher Brillouin shift when cultured in 2D environment than when cultured in 3D (0.0559 ± 0.023 GHz). Moreover, cells of the same type B also adopted a higher shift in the presence of the fibrillar topography than in the homogeneous 3D environment (-0.0333 ± 0.0131 GHz).

**Figure 2.**
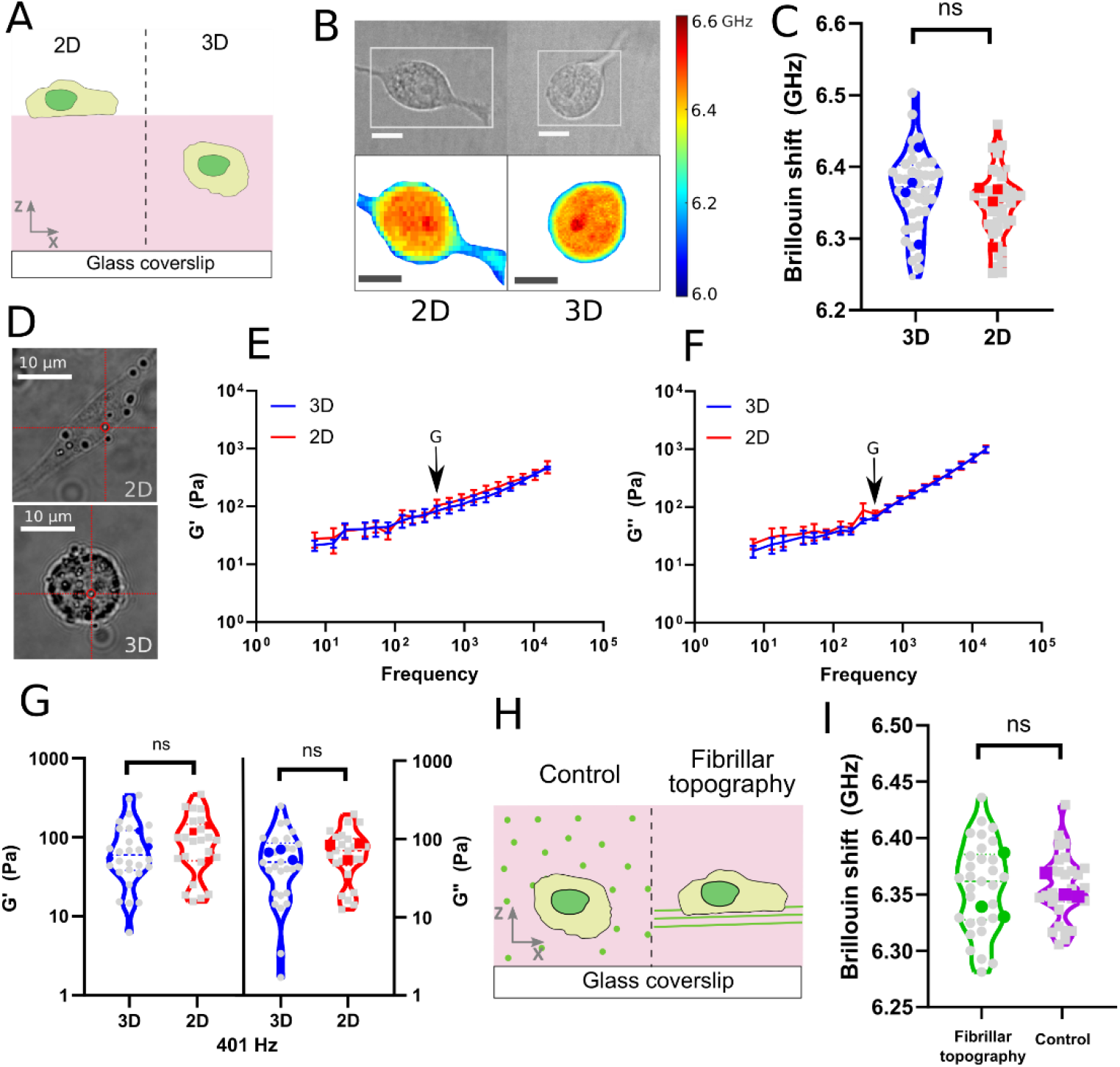
(A) Illustration of the two geometries in which we cultured U87 cells: on top of bulk Matrigel – 2D, and inside bulk Matrigel – 3D. (B) Example confocal Brillouin maps of cells in 2D and 3D configurations. Top images are brightfield images. The white rectangles indicate the location of the x-y confocal slices in which Brillouin shift was mapped. (C) Brillouin shift of cells grown in 2D vs 3D conditions. Unpaired t-test was performed on the average values of independent experiments (color points). (D) Example brightfield images of U87 cells with internalized 1 µm diameter polystyrene beads used in the optical trap measurements in 2D and 3D. (E) Elastic component (G’) of the shear modulus measured by the optical trap as a function of frequency ranging from 9 Hz to 15 kHz of cells in 3D (blue) and 2D (red). The error bar is standard error of the mean of three independent experiments. (F) Viscous component (G’’) of the shear modulus measured by the optical trap as a function of frequency ranging from 9 Hz to 15 kHz of cells in 3D (blue) and 2D (red). (G) Example shear modulus (G’) and loss modulus (G’’) at a single frequency denoted by arrows in (D) and (F). Each gray point is average of all the beads probed inside a single cell. Each color point is the average of an independent experiment. Two-way ANOVA was used to compare the 2D and 3D the average values for G’ and G’’ across three independent experiments and all frequencies. (H) Illustration of the fibrillar topography experiment in which we cultured U87 cells in presence of the fibrillar topography (FT) and in presence of homogeneously dispersed fibronectin coated magnetic beads (FT control). (I) Brillouin shift of cells grown in FT and FT control conditions. Unpaired t-test was performed on the average values of independent experiments (color points).

**Figure 3.**
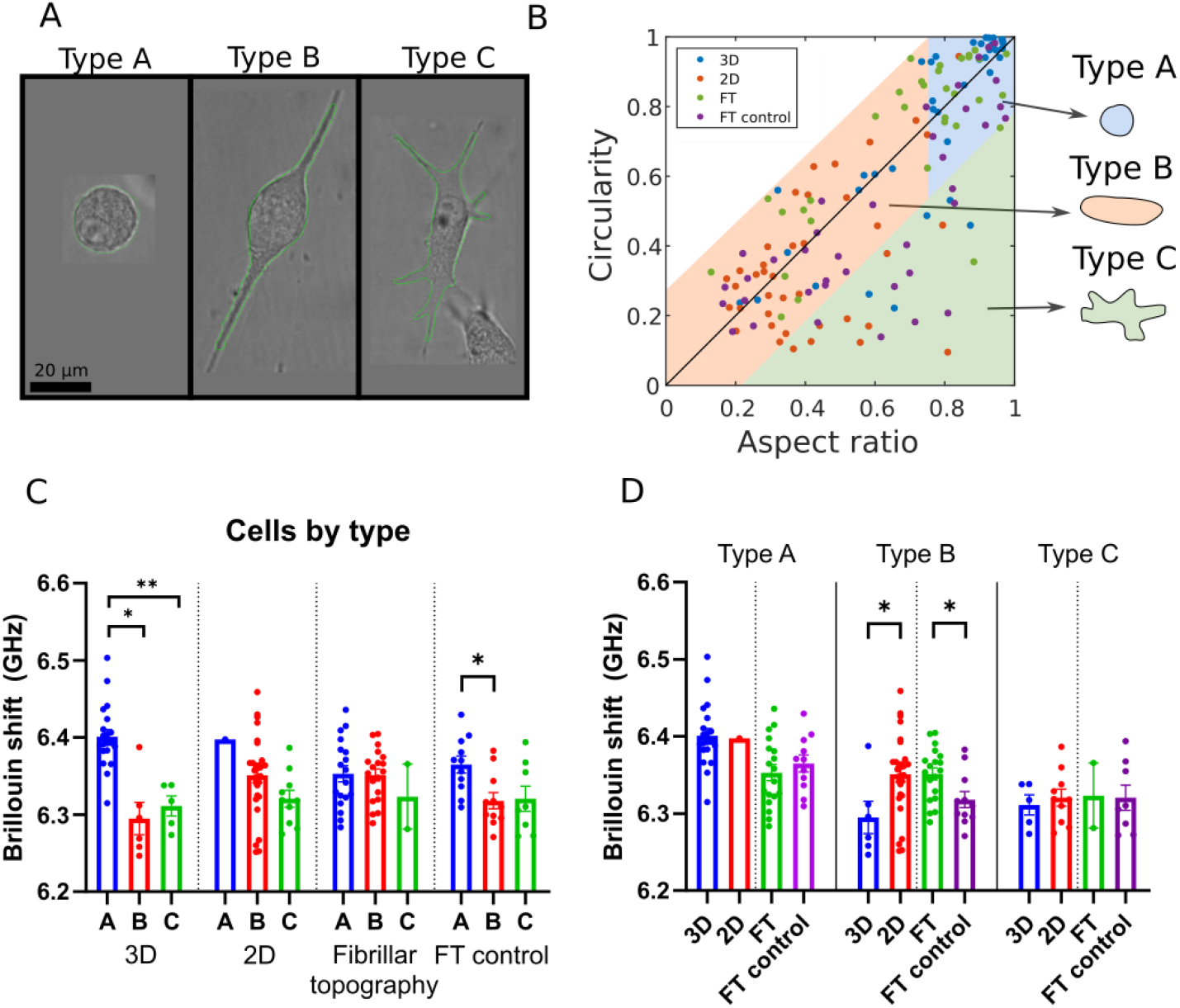
(A) Example brightfield images of U87 cells and the shape outlines (green line) (B) Scatter plot of circularity vs. aspect ratio (minor/major axis) of all cells. Cells were classified by type according to the region of the scatter plot in which they fall. Type C (irregular) cells are defined as cells that are more than 2 standard deviations away from the identity line. The rest of the cells is classified in type A and type B cells. Type A (round) cells are identified as cells that have aspect ratio larger than 0.75 (3/4). Type B cells (elongated) have aspect ratio less than (3/4). (C) Plot of average Brillouin shift of cells measured in the four conditions described in Fig 2., separated by cell type. Within some conditions there are differences in Brillouin shift of cells of different types. (D) Comparison of 3D vs. 2D and aligned vs. non-aligned Brillouin shift of all cells within a single cell morphology type. In type B group, Brillouin shift of 3D cells is lower than the Brillouin shift of 2D cells, while cells grown in presence of fibrillar topography (FT) have higher shift than the cells grown in an isotropic matrix (FT control). Unpaired t-test for comparing 3D versus 2D, and FT vs FT control conditions in each category with at least 3 cells.

**Figure 4.**
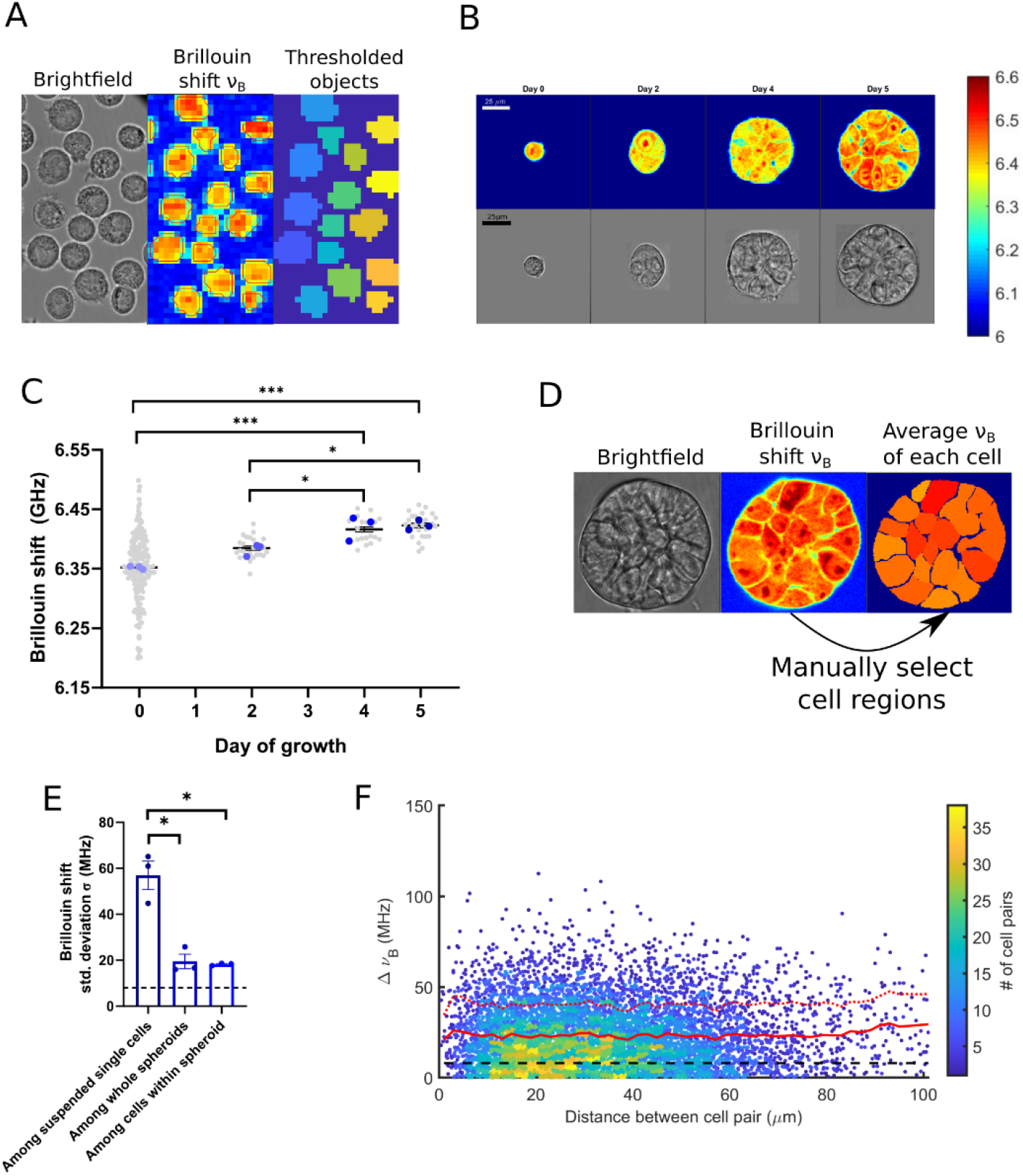
(A) Example Brillouin image and corresponding brightfield image of suspended cells in a dish. Individual cells were identified in the Brillouin shift images by thresholding and watershed transform (see Materials and Methods). (B). Example Brillouin shift maps of cells and spheroids at different stages of growth. (C) Average Brillouin shift of cells and whole spheroids (gray points) in three independent experiments (color points). Each experiment is average of multiple spheroids (n=7±2 spheroids), and in case of suspended cells (day 0) (n=58±3 cells). Statistical difference was detected using one-way ANOVA with Tukey correction performed on the average values of the three independent experiments (n=3). (D) Illustration of the process of identifying single cells within the spheroid. Single cells are manually selected in the Brillouin shift maps, and shift in each cell is averaged. (E) Standard deviation of measured Brillouin shifts of cells in three cases: across all single cells measured in an independent experiment (n=58±3 cells), across all spheroids in an independent experiment (n=7±2 spheroids), and across cells within a single spheroid (averaged over all spheroids in given experiment, n=17±6 cells, n_spheroids_=7±2). Dotted black line denotes instrumental precision of Brillouin shift measurements. (F) Absolute difference in Brillouin shift versus cell distance for all pairs of cells within single spheroids, across all day 5 spheroids of MCF10CA1h cells (n_pairs_=7266, Pearson correlation coefficient ρ=0.025). Color of scatter plot denotes the density of points. Distances were binned in the bins of size 2 µm and average shift (solid red line) and 1σ error (dashed red line) as a function of cell pair distance are plotted. Shift values at distances > 60 µm were smoothed with a 5-bin moving average to remove out the noise due to a small number of pairs in those bins.

### Cell mechanical properties in multicellular systems

To investigate cell mechanical properties in the context of 3D culture that includes cell-cell interactions, we turn to the *in vitro* method of 3D culture of breast epithelial cell spheroids. This method is well established and allows for repeatable formation of mammary gland like spheroids using laminin rich ECM (Matrigel). Here, we employed the MCF10CA1h (M3), a human breast cancer line. This cell line forms robust spheroids when cultured in laminin rich extracellular matrix (lrECM) using the “3D on-top” method [40, 41]. Using Brillouin microscopy, we mapped the mechanical properties of individual cells in suspension and longitudinally from day0 - day 5 during growth of spheroids. The heterogeneity of Brillouin shifts is smaller in spheroids when compared to single cells. The heterogeneity among spheroids is similar to the heterogeneity of single cells within those spheroids. To investigate the mechanical state of cells, we take advantage of the high-resolution nature of Brillouin microscopy to identify single cells in the Brillouin maps (Fig 4 E). By manually selecting each cell in each image, we can estimate the Brillouin shift of only the cells within the spheroid and thus remove effects of differential cell packing, or any cell-free spaces inside the spheroid. To check if the mechanical cooperativity is stronger between neighboring cells versus cells at the distant part of the spheroid, we looked at differences in Brillouin shift between each cell pair within a given spheroid. Plotting this data for all day 5 spheroids of M1 cells, we find that the average Brillouin shift difference for a pair of cells in a spheroid is constant function of pairwise distance between cells (Fig. 4 F).

## Discussion

Cancer cells exhibit distinct mechanical properties compared to those measured for the normal counterparts that are context dependent [2, 28, 66, 67]. One promising idea is to use mechanical phenotype to predict metastatic potential and drug responsiveness in order to aid diagnosis and treatment. However, the measured mechanical properties are determined by environmental factors such as availability and chemical identity of ligands [68], nearest neighbor interactions with cells [23], and ECM components [69]. In addition, the measured values of the cellular mechanical properties depend on the length and temporal scales at which the measurements are performed [26, 33, 70]. Thus, what is needed is the ability to measure mechanical properties of cells within native tissue microenvironments. Here, we employed a non-contact, label free technique, Brillouin microscopy to probe modulation of mechanical properties of cells in complex microenvironments. Furthermore, we employed broadband frequency range optical tweezers to probe the microrheology of cells to provide complementary sub-micron scale measurements to validate our Brillouin microscopy results in 3D complex *in vitro* system. We applied these techniques to established *in vitro* models which recapitulate *in vivo* basement membrane stoichiometry and tissue anisotropy. We find good agreement between two independent methods: optical trap and Brillouin microscopy for probing single cell mechanics. Here, we demonstrated that live cell-Brillouin microscopy revealed shifts that are sensitive to environmental conditions and pharmacological perturbations. We also longitudinally mapped mechanical changes as a function of spheroid formation from single cell to multicellular structures. We show that this technique is sensitive to heterogeneities that arise due to nearest neighbor effects and due to differences in cell morphology.

Mechanical phenotyping of tissues at multiple length scales have been of great interest in the field of mechanobiology. Optical based techniques such as optical stretchers [71], optical and magnetic tweezer-based modalities [33, 72, 73], real time deformation cytometry [74] have been employed to probe from nm scale to µm scale. However, many of these techniques for assessing material properties are unable to probe microscale mechanics for cells embedded in complex 3D tissues. We and others recently employed optical tweezer based active microrheology to map microscale mechanical properties in 3D culture models and in a living animal [33, 75-77]. Several technological advancements such as the *in-situ* trap stiffness calibration, and high sensitivity localization of the probe bead have enabled the applications in 3D cultures and in living animals [34]. Additionally, broad-band frequency analysis allows quantitation of distinct viscoelastic profile such as individual cell stiffnesses and hysteresivity, or relative liquid- or solid-like behavior. Access to the higher frequencies also can be used to probe additional rheological metrics such as a power law dependence in different materials. However, this and similar methods are limited to the use of introduced probes such as polystyrene beads or intrinsic components such as organelles to infer the underlying mechanical properties. In the case where external probes are introduced into tissue, these probes must be introduced in numbers and densities that do not compromise cellular integrity and function. This then results in under sampling of the cell interior at the level of single cells. In the latter case, the intrinsic organelles may be of different sizes and shapes which makes it difficult to translate underlying deformation of tissue to a rheological value. Moreover, the application of forces to these organelles may drive an unwanted perturbation in physiological function. Thus, a non-invasive method to map the mechanical properties of tissue with high resolution is desirable. Brillouin microscopy is an all-optical and purely non-invasive method. Thus, it provides an attractive and complementary approach to address this need.

Brillouin shift is a measure of the longitudinal elastic modulus, provided that we know the index of refraction and mass density of the material. In typical biological materials such as cells or tissues these parameters are not always known or directly measurable, and for that reason this and majority of other studies report values of Brillouin shift in GHz, rather than the value of the longitudinal modulus. Interestingly, for most cells and tissues the ratio of the refractive index and the square root of the density remains constant, since both index of refraction and density vary together, and the dominant contribution to Brillouin shift comes from the changes in M’. Therefore, Brillouin shift is used as a proxy measurement of the local longitudinal elastic modulus [44-46]. In most cells and tissues this assumption is valid, but there are exceptions like the lipid-rich samples where the mass density and index of refraction have dramatically different values [78, 79].

In contrast to the Young’s and shear moduli which are defined in terms of the specific engineering strains, longitudinal modulus is defined as uniaxial deformation due to uniaxial stress [43]. This elastic constant includes a volume change and therefore it is many orders of magnitude larger in incompressible materials. Furthermore, Brillouin scattering probes the longitudinal modulus on the GHz timescale [43], therefore the Brillouin derived longitudinal modulus is a fundamentally different elastic constant from the traditional measurements of Young’s modulus or shear modulus. However, we and others have previously shown empirical relationships between changes in the Brillouin shift and changes in the stiffness obtained using traditional rheological methods in cells and tissues, which have to be ascribed to a common dependence of the two moduli to underlying structural and biophysical factors [44, 80, 81]. Importantly, Brillouin scattering probes material mechanical properties with spatial scale of a few hundred nanometers which is determined by the characteristic length scale of the thermal density fluctuations [82, 83]; therefore, there is a mismatch between the length scales obtained between the Brillouin microscopy and AFM or bulk rheology measurements. Here, we experimentally compare Brillouin shift with a complimentary microrheology measurement at the same length scale using the optical trap method, which probes the material stiffness on the scale of ∼400 nm. However, there are two important differences between these methods. They measure different rheological parameters (shear versus longitudinal moduli), and they probe the mechanical properties at different timescales (Hz-kHz versus GHz). The observed good agreement of microscale properties between the two modalities at different timescales points to the fact that the underlying material structure of the live cell can be probed in different rheological experiments, and that cell’s mechanical phenotype determined in such a way can be used as a biomarker of the overall cell state.

Cells adopt different morphologies in different tissue environments [36, 84]. It has been postulated that cell morphology is linked to cell fate and may also be linked to predicting metastatic potential [65]. These studies have implicated physical traits such as cell morphology, mechanical phenotype, and migration as determinants of aggressiveness of many types of cancers such as breast, pancreatic, osteosarcomas, and prostate cancers [85-87]. These studies have also performed combinatorial morphometrics and transcriptomics in efforts to yield predictions of metastasis [60, 65, 88]. Differences in cell morphology point to the differences of overall cell state which may also reflect differences in mechanical properties [20, 44, 66, 89]. Hence, here we compared the cell mechanics as a function the environment as a function of the cell morphology. Here, we determined that for similar morphologies, cells adopt different mechanical phenotypes due to environmental conditions. Our findings support the idea that physical parameters such as the mechanical properties can also be a proxy for cell state in a context dependent manner. However, this and other data suggest that combinatorial analysis of multiple physical parameters such as morphology and mechanical properties will be needed to increase predictive capacity of cell fate and function.

Tissues are multicellular units that can be homogenous cell types or combinatorial organization of different types of cells. Organoid and 3D culture models allow the recapitulation of microtissues observed *in vivo* for a multitude of epithelial organs [25, 41, 90-93]. Moreover, they can also recapitulate the progressive disordered architectures associated with the malignant transformation [94, 95]. Dynamic cell behaviors such as coherent rotation [96], cell-cell adhesion turnover [23] and coordinated cell-ECM interactions [5, 6, 97] are important in the establishment of glandular tissues and become dysregulated in the malignant transformation. However, it is less understood how cells regulate crosstalk across multiple spatial and temporal scales. One idea is that there is a mechanical coupling that acts a feedback mechanism to facilitate multicellular growth. We recently determined that there is a microscale coupling between normal cells and matched organ ECM microenvironment using optical trap based active microrheology [33]. We also determined that there was a mismatch between intracellular cell mechanics with that of the surrounding matrix environment that was dependent on chemical specificity of the ECM hydrogel. This technique can be employed for longitudinal mapping of multicellular spheroids. However, in case of determination of multi-cellular coupling, the measurement throughput is limited as it heavily relies on the optimized introduction of external probes in every cell within a spheroid. Here, Brillouin microscopy rapidly determined the mechanical properties of individual cells and reveals co-operative behavior during the establishment of spheroids across multiple days. For single cells in 2D and 3D, cell shape in 2D brightfield images is a good approximation of the real cell shape. On the other hand, in spheroids cells are tightly packed and take complex shapes. In confocal Brillouin slices, cell boundaries can be easily distinguished, and we take advantage of this to approximate the Brillouin shift of each cell by manually selecting them. Note that, without imaging a full volume of the spheroid which is still impractical due to limited speed of acquisition (∼10-15 min per confocal slice of the spheroid), we cannot get an estimate of the complex cell shapes inside the spheroids. For this reason, we quantify the variance of average Brillouin shifts on the level of individual cells. The decreased heterogeneity among cells in spheroids points to the existence of mechanical cooperativity between cells that make up a single spheroid. It is important to note here, that the shift heterogeneity within spheroids is still larger than the instrumental precision of Brillouin shift, which means that we are still sable to detect cell-to-cell variability (∼20 MHz) which is smaller than the variability of single cells (∼45 MHz). Furthermore, we find that this variance is spatially constant across the spheroid as the differences in the mechanical properties of any two cells do not depend on their distance within the spheroid. It is important to note that the cell colonies start growing from single cells, and the uniformity in mechanical properties could be explained by their clonal nature.

In summary, using all-optical methods we quantify the mechanical properties of single cells and multicellular structures in 3D environments that were not accessible before. In 3D spheroids, we observed a lower variance of Brillouin shifts as compared to single cells. In addition, we have shown that the mechanical state of the cell is dynamic and changes depending on the context. In the future, it will be interesting to investigate the relationship between the expression of key cell-cell and cell-ECM interaction proteins in the spheroids in relation to the mechanical heterogeneity of cells, to elucidate the role of mechanical coupling of cells in multicellular systems.

## Acknowledgements

We thank Dr. Woong Young So for the help with the optical tweezer setup and experiments. We also thank Dr. Colin Paul for the assistance with the 3D fibrillar architecture microenvironment engineering. This effort was supported by the Intramural Research Program of the National Institutes of Health, the National Cancer Institute, NCI-UMD Partnership for Integrative Cancer Research, National Institutes of Health (R21-CA258008); National Science Foundation (CMMI 1929412).

## Author contributions

K.T., G.S., and M.N. designed the research. M.N. performed the research and analysis. K.T., G.S., and M.N. wrote the manuscript.

## Data availability

The raw data required to reproduce these findings are available from Kandice Tanner, Ph.D., 37 Convent Dr., Bethesda, MD 20852. The processed data required to reproduce these findings are available from Kandice Tanner, Ph.D., 37 Convent Dr., Bethesda, MD 20852.

